# MOTIFS IN SARS-COV-2 EVOLUTION

**DOI:** 10.1101/2023.01.27.525936

**Authors:** Christopher Barrett, Andrei C. Bura, Qijun He, Fenix W. Huang, Thomas J. X. Li, Christian M. Reidys

**Author notes:** Author contributions: C.B. and C.M.R. designed research; F.W.H. and T.J.X.L. collected data; A.C.B., Q.H., F.W.H., T.J.X.L. and C.M.R. performed research and analysis; C.B., A.C.B., F.W.H., Q.H., T.J.X.L. and C.M.R. wrote the paper.

## Abstract

We present a novel framework enhancing the prediction of whether novel lineage poses the threat of eventually dominating the viral population. The framework is based purely on genomic sequence data, without requiring prior established biological analysis. Its building blocks are sets of co-evolving sites in the alignment (motifs), identified via co-evolutionary signals. The collection of such motifs forms a relational structure over the polymorphic sites. Motifs are constructed using distances quantifying the co-evolutionary coupling of pairs and manifest as co-evolving clusters of sites. We present an approach to genomic surveillance based on this notion of relational structure. Our system will issue an alert regarding a lineage, based on its contribution to drastic changes in the relational structure. We then conduct a comprehensive retrospective analysis of the COVID-19 pandemic based on SARS-CoV-2 genomic sequence data in GISAID from October 2020 to September 2022, across 21 lineages and 27 countries with weekly resolution. We investigate the performance of this surveillance system in terms of its accuracy, timeliness and robustness. Lastly, we study how well each lineage is classified by such a system.

## Introduction

Current viral genomic sequencing efforts facilitate epidemiological surveillance close to real time [6]. The challenge now becomes the efficient identification of variants within these viral sequences that can pose possible threats. We present here a bottom up approach that can rapidly predict whether or not, at a given geographical location, a lineage will dominate the viral population. This assessment only takes as input a time series of multiple sequence alignments (MSAs) consisting of genomes sampled from the viral population up to the considered point.

The key idea is the identification of site motifs: maximal sets of polymorphic sites exhibiting co-evolutionary signals within the MSA, rather than considering the emergence of any one particular set of mutations. The sites that will manifest as motifs are selected on the basis of exhibiting sufficient mutational activity and satisfying a certain diversity criterion. Site motifs are then constructed using distances that can capture co-evolutionary coupling between site pairs. The collection of site motifs then represents the relational structure among the considered polymorphic sites.

Finally, we introduce a scoring function that evaluates the relevancy of a lineage, with respect to the relational structure present in the alignment. Using this score, we develop a genomic surveillance system that can predict whether a lineage will dominate the viral population at a given location and a given time. Namely, an alert will be issued with respect to a lineage at a given time-location pair if we observe significant changes of the score.

We then put this framework to the test, by analyzing retrospectively the sequence data obtained during the COVID-19 pandemic. Specifically, we consider SARS-CoV-2 genomics sequence data from October 2020 to September 2022, across 21 lineages and 27 countries with a weekly resolution. For the purpose of this analysis we shall derive notions of accuracy, timeliness and robustness and quantify its sensitivity and specificity in terms of classifying lineages at country resolutions as well as worldwide.

### Background

Genomic surveillance plays an instrumental role in combating rapidly mutating RNA viruses [17]. In particular, it was a vital necessity in the effective mitigation and containment of the COVID-19 pandemic [41, 10]. While mRNA vaccine development and distribution were successful in the US, the Omicron-variant, still gave rise to questions regarding their efficacy [8]. Timely vaccine development necessitates the rapid recognition of critical adaptations within SARS-CoV-2 [37, 27]. Genomic surveillance leverages applications of next-generation sequencing and phylogenetic methods to detect variants that are phenotypically or antigenetically different, facilitating early anticipation and effective mitigation of potential viral outbreaks [7].

One of the central tasks in genomic surveillance is the identification of virulent emerging variants, or of variants that have developed vaccine resistance. The designation of SARS-CoV-2 variants of concern/relevance exemplifies such an identification process [30, 48]. This designation is based on phylogenetic methods and involves four steps: lineage assignment, mutation extraction, biological analysis and declaration. First, a large phylogenetic tree is constructed from publicly available genomes and its sub-trees are examined and cross-referenced against epidemiological information to designate new lineages [3, 39]. Secondly, a collection of mutations frequently observed in a lineage, is extracted and defined to be characteristic for that lineage. The biological impact of this collection of mutations is then analyzed in wet-lab/in silico experiments. Thirdly, in the wake of identified biological features, such as an increase in transmissibility or severity, the lineage/variant is declared of concern/relevance.

Population-based approaches were developed to complement phylogenetic-based methods with the goal of rapidly identifying and monitoring critical mutations on the SARS-CoV-2 genome. Frequency analysis is widely used to monitor variant circulation [31, 35]. The increasing relative frequency of a mutation might indicate the emergence of a new variant [50]. Entropy measurements, derived from nucleotide frequency, highlight nucleotide positions with high variation and facilitate the compact representation of SARS-CoV-2 variants [14].

Mutations on viral genomes do not always appear independently. For example, the S:D614G change, caused by the nucleotide mutation A23404G on the SARS-CoV-2 genome, was almost always accompanied by three other nucleotide mutations: C241T, C3037T and C14408T [36]: these four positions exhibited a co-evolutionary pattern [50, 2]. In fact, it is by construction that positions in a molecule that share a common constraint do not evolve independently, and therefore leave a signature in patterns of homologous sequences [12, 38]. Extracting such co-evolutionary signals from a sequence alignment leads to a deeper understanding of the impact of mutations on viral fitness and can facilitate the early detection of emerging variants.

Currently, co-evolution detection strategies are based on observing the frequency of nucleotide combinations in two distinguished positions [33, 47]. In protein folding, MSAs are employed to identify related positions via mutual information [45]. While there are contributions on the level of networks: for instance [1] studies mutual information networks of enzymatic families in protein structures to unveil functional features, the work is focused on how to account for the effect of phylogeny on this identification [4]. To this end, two modifications to mutual information are introduced: row-column weighting [18] and average product correction [11]. Mutual information has also been used to detect coevolutionary signals in alignments of RNA sequences [21]. In [20, 49] statistical methods could differentiate correlation patterns induced by functional constraints from those induced by shared ancestry. These methods are concerned with pairwise relations since their objective is to determine RNA secondary structure. Thus, the idea of considering pairwise relations between columns within an MSA has been successfully used in protein and RNA folding more than a decade ago. Later we shall see that our notions of site motifs and relational structures represent an extension of this methodology, encapsulating *k*-ary relations present in the MSA.

### Technical overview

Co-occurring mutations exhibit traces of evolutionary selection pressure and fitness gained through synchronized mutations can be more significant than the total fitness when each mutation occurs independently [29, 25, 13]. Therefore, a collection of synchronized polymorphic sites is a result of selection pressure and represents a signal for selective advantage, worth incorporating in viral surveillance frameworks.

Let 𝒜 be a viral population consisting of sequences with *n* genomic sites (labelled as [*n*] := {1, …, *n*}). A *relational structure* on the population 𝒜, denoted by *S*_𝒜_ = {*M*_*1*_, …, *M*_*k*_}, is a collection of *site motifs*, or motifs for short, where each motif *M*_*i*_ ⊂ [*n*] consists of polymorphic sites that exhibit a similar evolutionary pattern in 𝒜, see Figure 1. Given an MSA 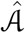 that consists of sampled sequences from 𝒜, we can approximate *S*_𝒜_ with 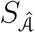. We detail the derivation of a relational structure from a given MSA in the Materials and Methods section.

**Figure 1.**
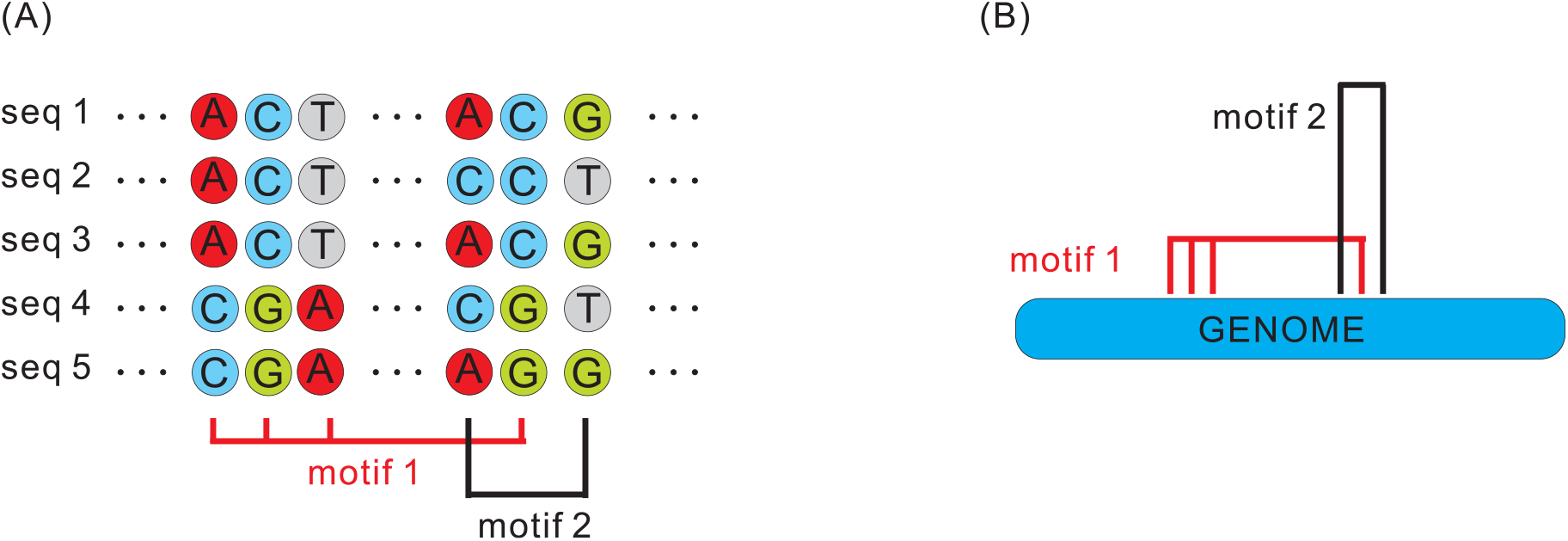
Co-evolutionary pattern in an MSA (A) and the induced relational structure (B).

The relational structure of a viral population is by construction not fixed and evolves over time, due to changes in selection pressure. Drastic changes in the relational structure, as we shall see, will be indicative of watershed events in viral such as the emergence of a new and fitter lineage. In the following, we shall illustrate the concept, via a case study, demonstrating the evolution of relational structures in the context of viral strain competition.

### Case study: evolution of the relational structure SARS-CoV-2 population in UK

We consider the strain competition in the UK SARS-CoV-2 viral population, from October week 3, 2020, to October week 4, 2021. During this time frame, the Alpha variant (lineage B.1.1.7) emerged, and quickly became the dominant strain in terms of relative frequency. After a period of time, the Alpha variant was then superseded by the Delta variant (lineage B.1.617.2), together with its AY sub-lineages (B.1.617.2+AY.*), see Figure 2.

**Figure 2.**
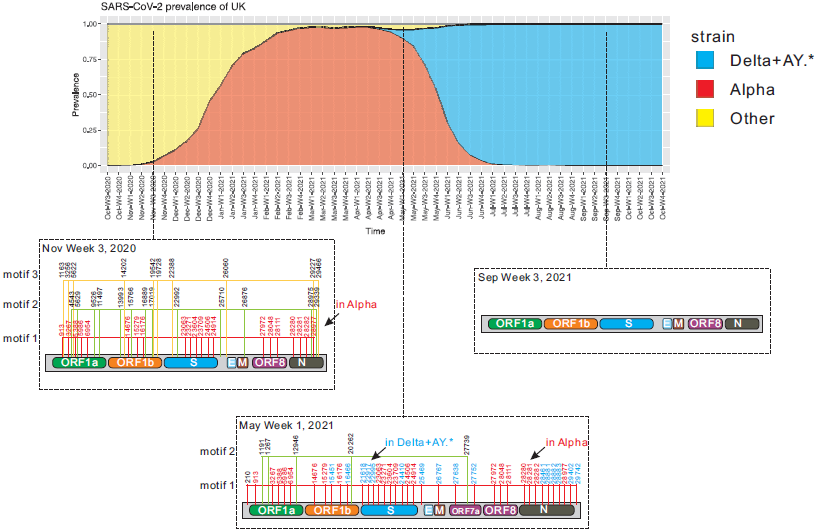
Evolution of the relational structure of the SARS-CoV-2 genome in the UK. (A) the relative frequency of key lineages (Alpha and Delta+AY.*.) from October week 3, 2020, to October week 4, 2021, in the UK. (B) the relational structure of SARS-CoV-2 at: 1) November week 3, 2020, when the relative frequency of Alpha is < 5%. 2) May week 1, 2021, when Delta+AY.* emerges and competes with Alpha. 3) September week 3, 2021, when the relative frequency of Delta+AY.* is > 95%. This analysis only considers relational structures with a minimum motif size of 5 and we only display the three largest motifs.

In Figure 2, we display the relational structures at three distinct moments in time: November, week 3, 2020 (emergence of Alpha). 2) May week 1, 2021 (emergence of B.1.617.2 (Delta) + AY.*). 3) September week 3, 2021 (domination of B.1.617.2 (Delta) + AY.*). In November week 3, 2020, the largest motif in the relational structure consisted of 21 sites. In fact, all of these 21 sites corresponded to characteristic mutations of Alpha. At this time, the relative frequency of Alpha is below 5%. The majority of the sequences in the populations carry the same nucleotide type as the original reference sequence on these 21 site. However, even at low relative frequency, the characteristic mutations on these 21 sites appear in a highly coordinated fashion. As a result, these sites exhibit a similar evolutionary pattern and form a motif, see Figure 2.

In May week 1, 2021, the largest motif in the relational structure consists of 38 sites. Among these 38 sites, 21 of them are precisely the 21 sites mentioned previously, that correspond to characteristic mutations of Alpha, while 16 of the remaining 17 sites correspond to characteristic mutations of the Delta+AY.* variants. At this time, the relative frequency of Alpha is above 90% while that of Delta+AY.* is below 5%. Note that on the 21 Alpha characteristic mutation sites, sequences in the Delta+AY.* variants are carrying the same nucleotides as the original reference sequence. On the other hand, sequences in the Alpha variants do not carry the 16 Delta+AY.* characteristic mutations. As such, the strain competition between Alpha and Delta+AY.* causes the formation of this large motif, see Figure 2.

In September week 3, 2021, Delta+AY.* becomes the dominant strain in the UK. The relative frequency of Delta+AY.* is so high, that we observe little diversity in the columns of the MSA and no motifs in the relational structure.

The purpose of this paper is to provide a meaningful augmentation to existing genomic surveillance methodology. The novelty of the proposed approach lies in focusing on the linkage and not the specific patterns of mutations appearing in the data. This leads to the concept of motifs and relational structures. The proposed framework is a stepping stone to a more systematic development of the notion of relational structures for general aligned data.

## Results

In this section, we present an approach based on relational structures in order to enhance genomic surveillance. Our system will issue an alert regarding a lineage, based on its effect on the relational structure. We will conduct a comprehensive retrospective analysis and investigate the performance of this surveillance system studying its accuracy, timeliness and robustness. Finally, we will investigate how well each lineage is classified by our system.

### Genomic surveillance based on relational structures

We introduce a novel measurement *ε*(*L, S*) (see Materials and Methods), that tracks the relevancy of a lineage *L* with respect to a relational structure *S*. The time evolution of *ε*(*L, S*(*t*)) can provide insights into the dynamics of a lineage *L* within the viral population. We develop a surveillance system based on the difference function Δ*ε*(*L, S*(*t*)) = *ε*(*L, S*(*t*)) − *ε*(*L, S*(*t* − 1)). We then perform a comprehensive retrospective analysis of the COVID-19 pandemic as well as of the performance of the surveillance system in predicting lineage development.

We investigate the development of 21 SARS-CoV-2 lineages in 27 different countries with a weekly resolution, from October Week 3 2020 to September Week 4 2022 (94 weeks in total). The lineages we consider are selected among those monitored by the BV-BRC SARS-CoV-2 tracking system [5, 34], see Table 1. Since shared mutations among lineages transcend into similarities of their corresponding motifs, lineages are selected such that there is no substantial overlap of their characteristic mutations, see the Materials and Methods section for details of this selection procedure.

**Table 1.** Selected lineages.

| Lineage | WHO name | Emergence location | Emergence time |
| --- | --- | --- | --- |
| B.1.617.2+AY.* | Delta | India | October 2020 |
| B.1.1.529+BA.* | Omicron | Botswana | November 2021 |
| B.1.1.138 | - | UK | February 2021 |
| B.1.1.519 | - | Mexico | October 2020 |
| B.1.1.523 | - | Multiple | August 2020 |
| B.1.1.7 | Alpha | UK | September 2020 |
| B.1.351 | Beta | South Africa | May 2020 |
| B.1.427 | Epsilon | USA | September 2020 |
| B.1.466.2 | - | Indonesia | August 2020 |
| B.1.525 | Eta | Multiple | December 2020 |
| B.1.526 | Iota | USA | November 2020 |
| B.1.617.1 | Kappa | India | October 2020 |
| B.1.617.3 | - | India | October 2020 |
| B.1.619.1 | - | Multiple | May 2020 |
| B.1.620 | - | Multiple | November 2020 |
| B.1.621 | Mu | Columbia | January 2021 |
| C.1.2 | - | South Africa | May 2021 |
| C.37 | Lambda | Peru | December 2022 |
| P.1 | Gamma | Brazil | November 2020 |
| P.2 | Zeta | Brazil | January 2021 |
| R.1 | - | Multiple | October 2020 |

The countries examined were selected by continent: we picked the top five countries from each continent with respect to their number of contributed sequences to the GISAID [43] database, while for Oceania we only tracked the top two, see Table 2.

**Table 2.** Alert summary for each country, integrated over all lineages.

| Country | Continent | total | relevant | irrelevant | accuracy |
| --- | --- | --- | --- | --- | --- |
| USA | North America | 6 | 3 | 3 | 50.0% |
| Panama | North America | 7 | 3 | 4 | 42.9% |
| Mexico | North America | 9 | 4 | 5 | 44.4% |
| Canada | North America | 6 | 3 | 3 | 50.0% |
| Costa Rica | North America | 8 | 7 | 1 | 87.5% |
| UK | Europe | 3 | 3 | 0 | 100.0% |
| Sweden | Europe | 3 | 3 | 0 | 100.0% |
| Germany | Europe | 2 | 2 | 0 | 100.0% |
| France | Europe | 4 | 4 | 0 | 100.0% |
| Denmark | Europe | 4 | 4 | 0 | 100.0% |
| South Korea | Asia | 3 | 1 | 2 | 33.3% |
| Japan | Asia | 9 | 7 | 2 | 77.8% |
| Israel | Asia | 10 | 8 | 2 | 80.0% |
| Indonesia | Asia | 7 | 5 | 2 | 71.4% |
| India | Asia | 15 | 6 | 9 | 40.0% |
| Peru | South America | 9 | 6 | 3 | 66.7% |
| Colombia | South America | 12 | 4 | 8 | 33.3% |
| Chile | South America | 10 | 5 | 5 | 50.0% |
| Brazil | South America | 5 | 4 | 1 | 80.0% |
| Argentina | South America | 13 | 4 | 9 | 30.8% |
| South Africa | Africa | 25 | 7 | 18 | 28.0% |
| Nigeria | Africa | 16 | 8 | 8 | 50.0% |
| Mauritius | Africa | 12 | 6 | 6 | 50.0% |
| Kenya | Africa | 5 | 4 | 1 | 80.0% |
| Botswana | Africa | 16 | 11 | 5 | 68.8% |
| New Zealand | Oceania | 11 | 7 | 4 | 63.6% |
| Australia | Oceania | 8 | 7 | 1 | 87.5% |

We denote a lineage as a *key lineage* for a given country, if its relative frequency within the viral population of that country reaches 50% at some point within our 94 week time window. Note that the notion of key lineage is by construction country dependent.

We now describe our surveillance system based on relational structures. Its aim is to predict whether a lineage *L* will develop into a key lineage for a given country *X*. with respect to *X*, an alert will be issued by our system, regarding lineage *L* at time *t*, if Δ*ε*(*L, S*_*X*_(*t*)) *>* 0.25, where *S*_*X*_(*t*) is the relational structure corresponding to *X* at time *t*.

Note that a lineage can trigger, for the same country, multiple alerts over a period of time. A lineage can also trigger multiple alerts across multiple countries over the same period of time.

An alert with respect to a lineage *L*, in a given country, will be labelled as *relevant*, if the relative frequency of *L* reaches 50% within the next three months in said country. Otherwise, the alert will be labelled *irrelevant*.

### Accuracy

Integrated overall 27 countries and 21 lineages, our surveillance system issued 238 alerts in total. Of those, 136 were labelled as relevant (i.e. a 57.1% accuracy). The accuracy of the alerts for all lineage country combinations is displayed in Figure 3.

**Figure 3.**
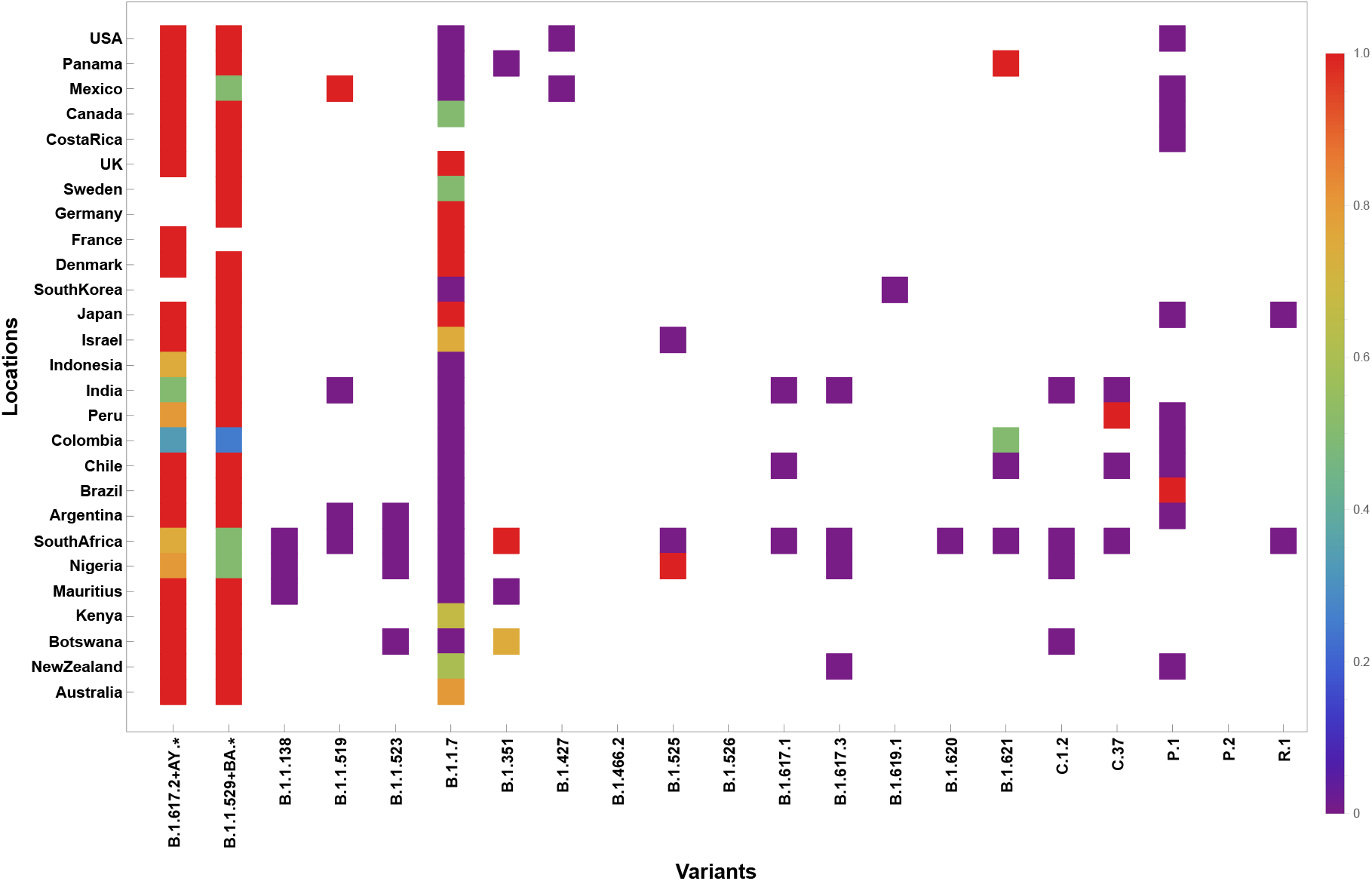
Accuracy of alerts across different variants and geographic locations. The x-axis denotes the variants considered, while the y-axis denotes the countries. The color of each dot represents the accuracy of our alert system with respect to the particular variant at the corresponding location. An empty cell indicates no alert is issued for that country lineage pair.

For the vast majority of the lineage-country combinations (437 out of 567 possible pairs), our system never issues an alert during the selected time span. Lineages B.1.1.7 (Alpha), B.1.617.2+AY.* (Delta) and B.1.1.529+BA.* (Omicron) are well-known for circulating globally [16, 44, 46, 19]. These three lineages received alerts from our system in almost all countries and the majority of these alerts were accurate (with the exception of some alerts for Alpha in certain countries). For lineages that circulate locally, our system still accurately predicts their dominance in certain specific countries, for instance, B.1.1.519 in Mexico, B.1.351 in South Africa and B.1.525 in Nigeria, while it remains silent in most other countries.

We proceed by investigating alerts issued for each specific country, integrated over all lineages, see Table 2.

The number of alerts varies across countries, with a mean of 8.8 and a standard deviation of 5.2. Germany has only 2 alerts in total, while South Africa has 25. The alert accuracy also varies: all counties in Europe exhibit an accuracy of 100.0% while those in South Africa have only 28.0% accuracy. It is worth mentioning that geographically adjacent countries tend to have similar alert patterns, which could be due to their viral populations having similar dynamics.

Finally, we investigate the alerts issued with respect to specific lineages, integrated across all countries, see Table 3.

**Table 3.** Alert summary for each lineage, integrated over all countries.

| Lineage | total | relevant | irrelevant | accuracy |
| --- | --- | --- | --- | --- |
| B.1.617.2+AY.* | 71 | 63 | 8 | 88.7% |
| B.1.1.529+BA.* | 35 | 29 | 6 | 82.8% |
| B.1.1.138 | 4 | 0 | 4 | 0.0% |
| B.1.1.519 | 6 | 2 | 4 | 33.3% |
| B.1.1.523 | 5 | 0 | 5 | 0.0% |
| B.1.1.7 | 56 | 22 | 34 | 39.3% |
| B.1.351 | 9 | 5 | 4 | 55.6% |
| B.1.427 | 3 | 0 | 3 | 0.0% |
| B.1.466.2 | 0 | 0 | 0 | NA |
| B.1.525 | 4 | 2 | 2 | 50.0% |
| B.1.526 | 0 | 0 | 0 | NA |
| B.1.617.1 | 3 | 0 | 3 | 0.0% |
| B.1.617.3 | 4 | 0 | 4 | 0.0% |
| B.1.619.1 | 1 | 0 | 1 | 0.0% |
| B.1.620 | 1 | 0 | 1 | 0.0% |
| B.1.621 | 5 | 2 | 3 | 40.0% |
| C.1.2 | 6 | 0 | 6 | 0.0% |
| C.37 | 5 | 1 | 4 | 20.0% |
| P.1 | 18 | 2 | 16 | 11.1% |
| P.2 | 0 | 0 | 0 | NA |
| R.1 | 2 | 0 | 2 | 0.0% |

The number of alerts varies significantly by lineages, with a mean of 11.3 and a standard deviation of 19.1, with 68.1% of alerts being issued to B.1.1.7 (Alpha), B.1.617.2+AY.* (Delta) and B.1.1.529+BA.* (Omicron) (which were all circulating globally at their respective times). Alerts issued to B.1.617.2+AY.* have the highest accuracy (88.7%). Finally, while alerts issued with respect to some lineages have 0% accuracy, they are sporadic and are generally issued with the “irrelevant” designation.

### Timeliness

For an alert system to be useful, one would expect that relevant alerts are issued at times when their corresponding lineages exhibit low relative frequency. As such, we investigate the relative frequency of lineages that issue relevant alerts.

If there exist multiple relevant alerts for the same lineage in a given country, we consider only the first issued relevant alert for that lineage. Integrating over all countries, there are a total of 69 relevant “first” alerts. For each such alert, we investigate the relative frequency of the associated lineage in the week prior to the issuance of the alert. If the relative frequency is below 5%, we consider the alert to be *timely*. Of the 69 relevant “first” alerts, 42 (60.9%) are timely. The ratio of timely alerts versus “first alerts” for each country is displayed in Figure 4, while the same for each lineage is shown in Figure 5.

**Figure 4.**
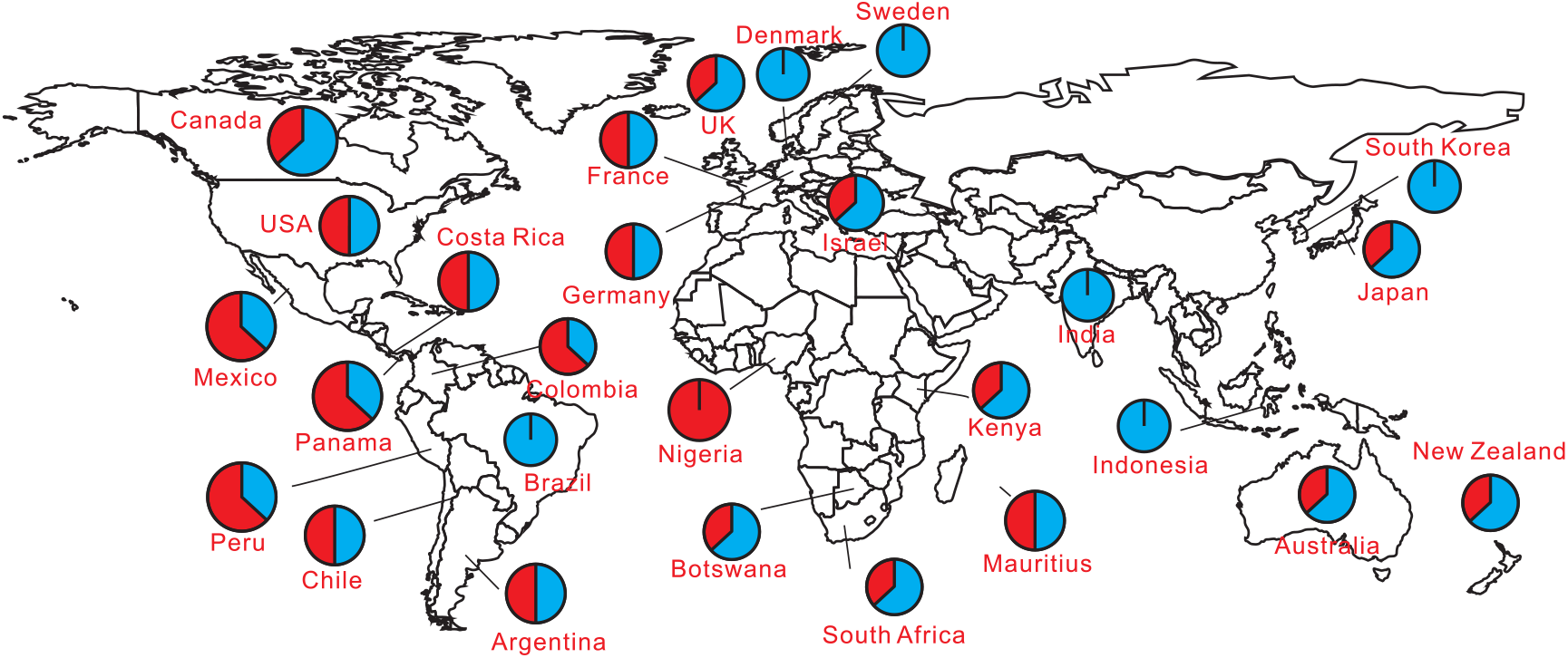
Timeliness of alerts across the globe. We illustrate the portion of timely first alerts (blue) versus non-timely first alerts (red).

**Figure 5.**
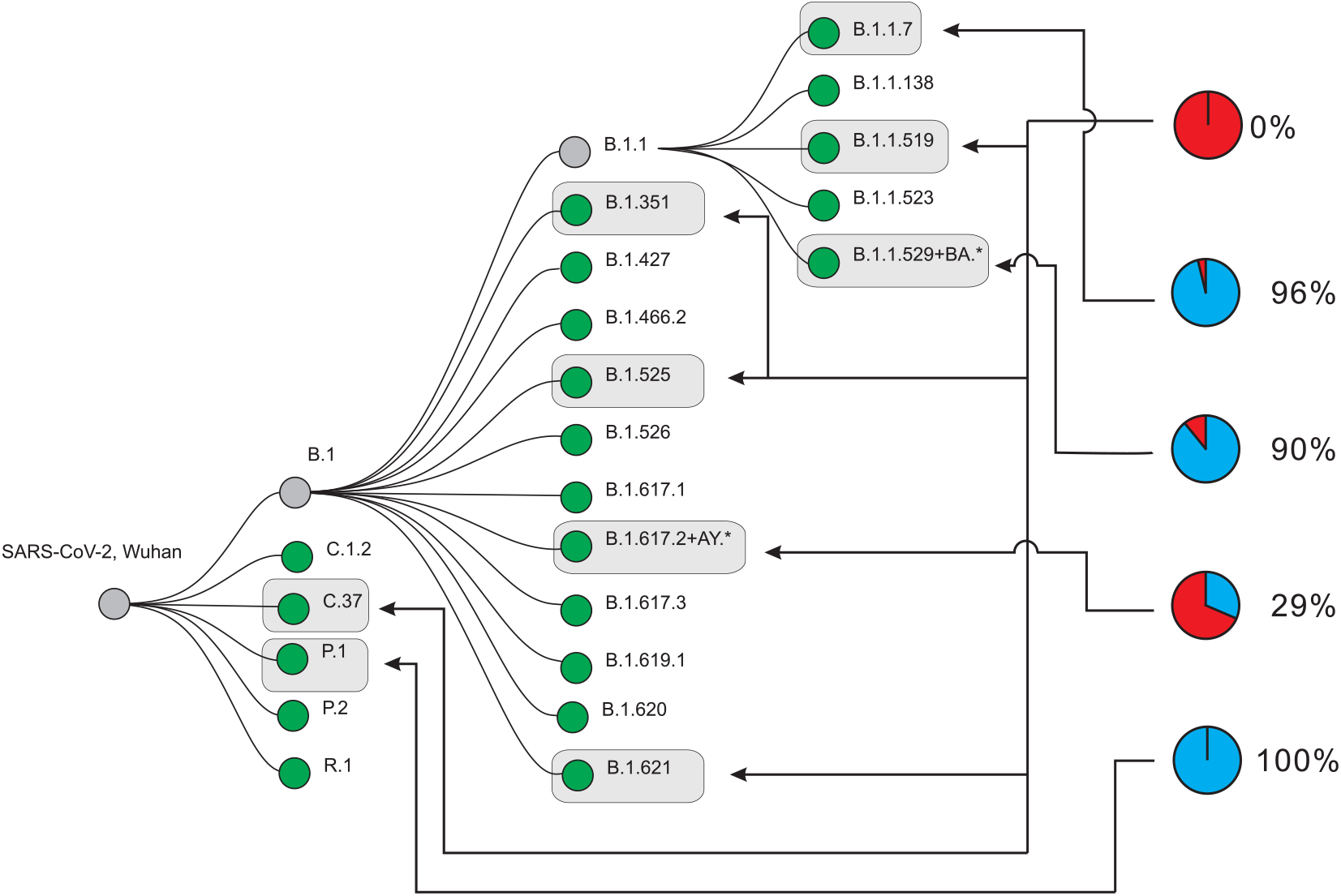
Timeliness of alerts across lineages. Lineages that have been issued at least one relevant alert over our study period (gray box). We illustrate the portion of timely first alerts (blue) versus non-timely first alerts (red) for each of these lineages. The associated percentage the ratio of timely first alerts to the total number of first alerts. The lineages are organized in a hierarchical tree structure according to their phylogenetic relations. Note that the 21 lineages considered in the study are lineages are somewhat independent in terms of overlaps of characteristic mutations

### Alert robustness

In the previous sections, we described various notions of effectiveness of the enhancement of viral surveillance based on relational structures. The alerts were derived from time-stamped MSAs, where each MSA consisted of sequences collected within a specific time period for a given country. However, sequencing capabilities across different countries are vastly different: in GISAID, the average number of sequences per week in Botswana during the investigative period is 48, while the average number of sequences per week in the UK is 22482. In view of this and in order to facilitate cross-country comparisons we study a notion of robustness of alerts.

To this end, we consider the robustness of the alerts for three lineages at three different time points, in the UK. We construct MSAs of different sizes via uniform subsampling with weekly sampling sizes of *M* = 50, 100, 500, 1000. For a given time point and fixed sampling size, we construct 100 pairs of MSAs, via sampling uniformly randomly from all UK sequences collected within that time frame. We then investigate the consistency in alert issuance behavior across the 100 pairs. For each sample size, the issuance of an alert can be considered as an indicator random variable *X* (*X* = 1 for issuance and *X* = 0, otherwise). Then, the variance of this random variable, 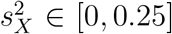, measures the inconsistency of the system’s output at that sample size, see Figure 6.

**Figure 6.**
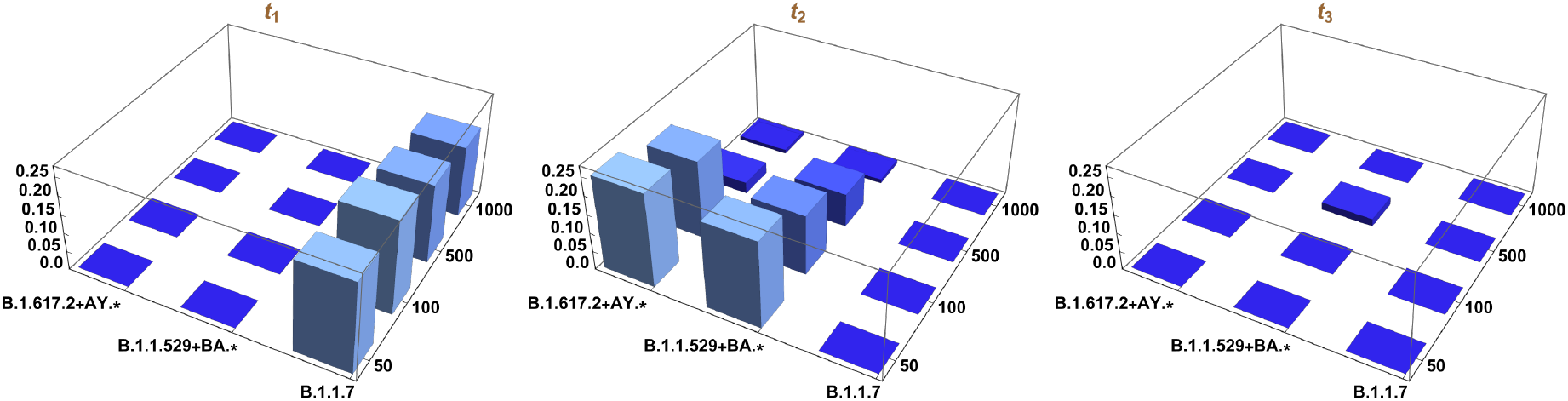
Alerts and robustness with respect to different sample sizes at three different times (*t*_*1*_: Nov week 4, 2020 to Dec week 1, 2020, *t*_*2*_: Nov week 4, 2021 to Dec week 1, 2021, *t*_*3*_: Jan week 2, 2022 to Jan week 3, 2022). In each figure, on the z-axis we display the variance of the random variable *X* (*X* = 1 for issuance and *X* = 0, otherwise) associated to a lineage-sample size pair.

Note that originally, our system issued alerts for B.1.1.7 at *t*_*1*_, and for B.1.617.2+AY.* and B.1.1.529+BA.* at *t*_*2*_ (this was initially achieved by sampling a single alignment of size 3000). The system did not issue alerts for the remaining time-lineage pairs. For the six combinations that correspond to this non-issuance, we observe that the system produces consistent output 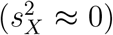, even at smaller sample sizes. In fact, the system almost faithfully reproduces the original non-issuance behavior. For the three combinations that correspond to alert issuance, we observe an increase in consistency as the sample size increases as measured by the variance. However, the rate of increase in consistency for the three combinations varies. For B.1.617.2+AY.* and B.1.1.529+BA.*, at *t*_*2*_ and sample size 500, the system reproduces the original issuance behavior more than 90% of the time, while the inconsistency for the alert issuance behavior associated to B.1.1.7 at *t*_*1*_ is still very high 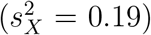, even at sample size 1000.

To state the obvious, it is critical to include all available sequences for surveillance purposes. The emergence of new variants is difficult to observe via small array-sizes. This being said: typically, 1000 sequences are sufficient to provide a robust alert. In the case of this study, the majority of countries attain this weekly sequencing rate. However when a lineage is at very low relative frequency and the strain composition is very heterogeneous then limited sampling is simply insufficient to provide a robust alert.

### Sensitivity and specificity

In the following, we shall shift focus from alerts to lineages and investigate whether or not they are correctly classified. Alerts as discussed previously are either relevant or not, while a lineage will be classified as true/false positive or true/false negative.

For a given country, we are interested in how many key lineages are successfully flagged and how many non-key lineages never trigger an alert. The first such number measures the sensitivity of the system while the second one measures the specificity. To this end, for a given country, denote by *P* the number of key lineages, and by *N* the number of non-key lineages. The number of key lineages that trigger relevant alerts in our system will be denoted by *T P*, while the number of key lineages that do not trigger relevant alerts will be denoted by *FN*. On the other hand, the number of non-key lineages that never issue an alert will be denoted by *T N*, while the number of non-key lineages that issue any alert will be denoted by *FP*. Sensitivity will then be given by 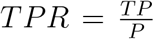, while specificity will be given by 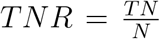. Integrated over all countries, we have *P* = 81 with *T P* = 69 and *T PR* = 85.2%. Furthermore, we have *N* = 486, with *T N* = 429 and *T NR* = 88.3%. See Figure 7 for an illustration of the relation between these quantities. Sensitivity and specificity for each individual country are displayed in Table 4.

**Table 4.** Country specific sensitivity and specificity for the alert system.

| Country | P | N | TP | TPR | TN | TNR |
| --- | --- | --- | --- | --- | --- | --- |
| USA | 3 | 18 | 2 | 66.7% | 16 | 88.9% |
| Panama | 3 | 18 | 3 | 100.0% | 16 | 88.9% |
| Mexico | 3 | 18 | 3 | 100.0% | 15 | 83.3% |
| Canada | 3 | 18 | 3 | 100.0% | 17 | 94.4% |
| CostaRica | 4 | 17 | 2 | 50.0% | 17 | 100.0% |
| UK | 3 | 18 | 3 | 100.0% | 18 | 100.0% |
| Sweden | 3 | 18 | 2 | 66.7% | 18 | 100.0% |
| Germany | 3 | 18 | 2 | 66.7% | 18 | 100.0% |
| France | 3 | 18 | 2 | 66.7% | 18 | 100.0% |
| Denmark | 3 | 18 | 3 | 100.0% | 18 | 100.0% |
| SouthKorea | 2 | 19 | 1 | 50.0% | 17 | 89.5% |
| Japan | 3 | 18 | 3 | 100.0% | 16 | 88.9% |
| Israel | 3 | 18 | 3 | 100.0% | 17 | 94.4% |
| Indonesia | 3 | 18 | 2 | 66.7% | 17 | 94.4% |
| India | 2 | 19 | 2 | 100.0% | 13 | 68.4% |
| Peru | 3 | 18 | 3 | 100.0% | 16 | 88.9% |
| Colombia | 3 | 18 | 3 | 100.0% | 16 | 88.9% |
| Chile | 3 | 18 | 2 | 66.7% | 14 | 77.8% |
| Brazil | 4 | 17 | 3 | 75.0% | 16 | 94.1% |
| Argentina | 2 | 19 | 2 | 100.0% | 15 | 78.9% |
| SouthAfrica | 3 | 18 | 3 | 100.0% | 6 | 33.3% |
| Nigeria | 3 | 18 | 3 | 100.0% | 13 | 72.2% |
| Mauritius | 3 | 18 | 2 | 66.7% | 16 | 88.9% |
| Kenya | 3 | 18 | 3 | 100.0% | 18 | 100.0% |
| Botswana | 3 | 18 | 3 | 100.0% | 15 | 83.3% |
| NewZealand | 3 | 18 | 3 | 100.0% | 16 | 88.9% |
| Australia | 4 | 17 | 3 | 75.0% | 17 | 100.0% |

**Figure 7.**
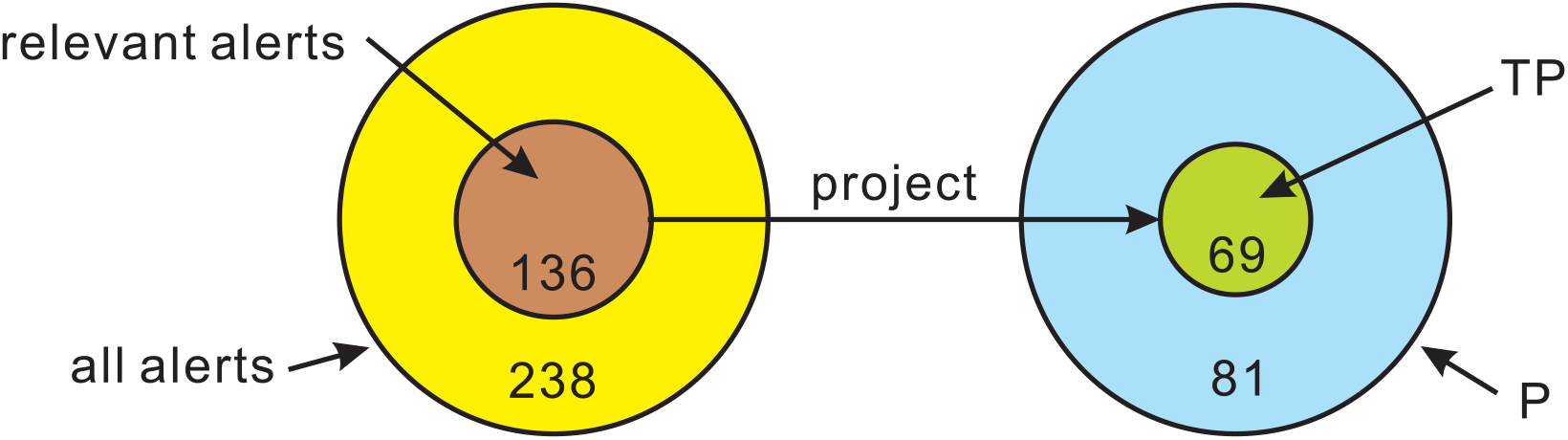
Accuracy versus Sensitivity. The system issues a total of 238 alerts, 136 of which being relevant, with an overall accuracy of 57.14%. We observe 81 key lineage instances (P) across all countries, 69 of which being issued at least one relevant alert (TP). The overall sensitivity of the system is 85.19%.

We note that *TPR* (sensitivity) and *TNR* (specificity) depend on the alert threshold *θ* of Δ*ε*(*L, S*_*X*_(*t*)). In general, if *θ* decreases, additional alerts will be issued and hence it is more likely that key lineages will be flagged by relevant alerts. This will lead to an increase in *T P* and hence an increase in sensitivity. However, at the same time, *T N* will decrease as it is more likely that non-key lineages will be flagged by alerts, which in turn reduces specificity. To quantify this trade-off, we perform a receiver operating characteristic (ROC) analysis [15]. Each *θ* induces a pair (*FPR, TPR*), where *FPR* = 1 − *T PR*. Varying *θ* ∈ [0, 1], produces the ROC curve [15], where the *x*-axis and *y*-axis represent the *FPR* and *TPR*, respectively. ROC curves for each of the countries considered, as well as a global ROC curve (integrated over all countries considered), are displayed in Figure 8.

**Figure 8.**
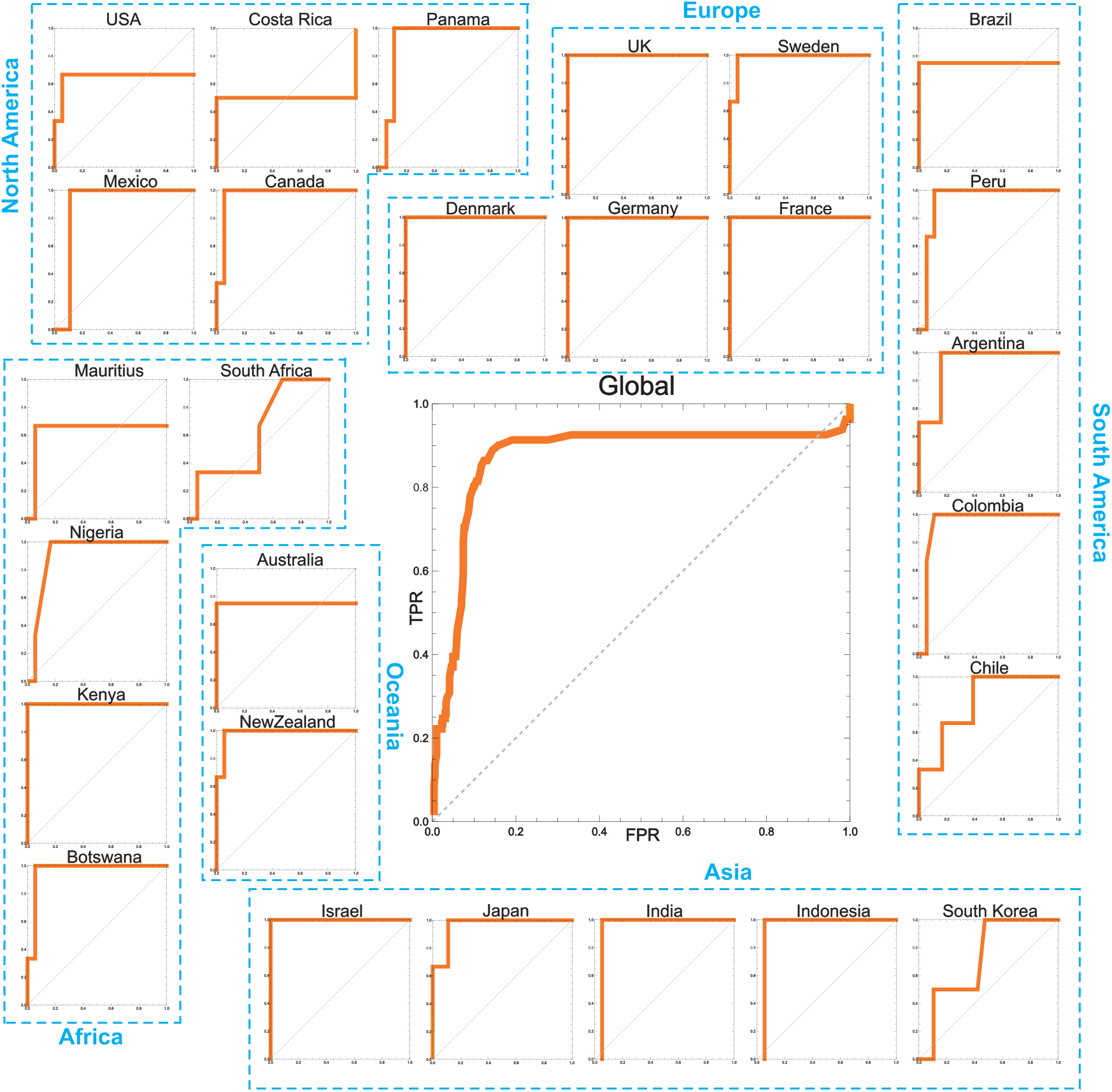
ROC curves across the globe. The *x*-axes and *y*-axes represent *FPR*s and *T PR*s, respectively.

## Discussion

In this paper we present an augmentation of genomic surveillance via a novel approach. The concept of motifs and relational structure is shown to provide at times accurate and timely predictions with respect to the development of viral lineages.

The key idea behind this approach is the study of site co-occurring mutational patterns that quantify MSA evolution. This is not new, however, what is novel is the focus not on the specific pattern but the relational linkage between sites. Genomic adaptation is, to a large extent, not facilitated by isolated mutations but rather by collections of mutations. Functionally connected mutations changing simultaneously therefore provide a key indicator of relevant evolutionary dynamics, and even at low frequencies, co-occurring mutations are indicative of the existence of noteworthy biological mechanisms inducing differentials in said dynamics.

The relational structure notion introduced here captures maximal sets of sites that simultaneously experience selection pressure. Differential changes of the relational structure within the MSA provide crucial information about the viral “heartbeat” since these are induced by selection: typically changes for the worse from the perspective of the viral host. We have shown that selection pressure leads to a small number of motifs that appear as distinguished patterns within the MSA and thus on the one hand, the method represents a significant reduction in data, and on the other hand, by identifying sites, it facilitates subsequent biological analysis.

The relational structure framework can be viewed as an orthogonal complement to current sequence-centric genomic surveillance. Phylogenetic analysis and lineage designation focus on the evolutionary closeness of genomic sequences while the relational structure encapsulates the interaction among sites in the MSA. As we demonstrated in the results section, combining the relational structure with lineage information, in the form of characteristic mutations, can provide us with deeper insights into how the lineage itself emerges and develops. Our score *ε*(*L, S*) captures by design the impact a lineage has on the formation of large motifs within the relational structure, and thus can provide information beyond characteristic mutation frequencies by capturing the degree of coordination between said characteristic mutations. A high degree of coordination provides a sensible statistical signal which can facilitate timely predictions.

One of the key challenges in combining the relational structure approach with lineage information is that they are in principle not always compatible. In our approach they are linked via the characteristic mutations corresponding to each lineage. For each lineage we obtained the characteristic mutation list from the Pango data base [39]. However currently, there are no generally agreed upon universal rules to define the characteristic mutations of a lineage. For a given lineage, the list of characteristic mutations can differ across different databases and each database has its own rules for defining characteristic mutations. Furthermore, characteristic mutations are not “characteristic” enough with the same mutation sometimes being used to characterize multiple lineages [24]. This is particularly the case, when one lineage is a sub-lineage of the other. To reduce the impact introduced by the hierarchical structure within the lineages themselves, for this paper we selected and merged lineages so that the sequences became better partitioned. In subsequent publications we will address the natural question that arises from this consideration, namely how to address hierarchical structures within designated lineages.

Currently there is no universal system that performs “well” for viral taxonomy below the level of species. In [40], a dynamic nomenclature approach was proposed for SARS-CoV-2 lineages based on phylogenetic methods. However, their approach still has the issue of over-designation, as more than 2500 lineage/sub-lineage names are assigned [39] and only a handful of them are considered to be critical from an epidemiological perspective. Our hope is that, by bringing relational structures into the picture, we can identify key lineages in a more accurate and timely fashion.

In fact, this approach can be generalized to a comprehensive system for classifying genetic diversity within a population and designating (sub)lineages. The key to such a general system is inferring the hierarchical (sub)lineage structure from the relational structures. More precisely, one could use each relational structure to construct a partition of the corresponding sequence population into non-overlapping sub-samples, that exhibit similar evolutionary patterns which support said relational structure. For each sub-sample, one then recursively computes its relational structure, which in turn yields (sub)lineages for the given sub-population. By this procedure, one could recover the hierarchical structure between lineages in an effective manner and without specialized human input, as is the case for Pango.

## Materials and Methods

### Lineage selection

We begin with all 28 lineages of concern that are currently or have been previously monitored by the BV-BRC SARS-CoV-2 tracking system [5, 34]. A lineage of concern (LOC) is a group of closely related viral sequences that exhibit similar mutations or recombinations that may significantly affect vaccine efficacy, transmissibility or disease outcomes [5]. The designation of LOCs is based on phylogenetic methods, and cross-referenced against epidemiological information [3, 40].

According to Nextstrain and CoVariants [22, 24], these lineages exhibit a hierarchical structure which reflects the evolutionary trajectories the viral population undertook when acted upon by selection pressures, see Fig. 9. For example, BA.1 and BA.2 are sub-lineages of Omicron; BA.4 and BA.5 are sub-lineages of BA.2. Such a hierarchical structure affects the lineage assignment of viral genomic sequences, as sequences in a sub-lineage are also assigned to their ancestors. This lineage assignment method has been adopted by the CDC during their genomic surveillance efforts [6]. However, this is not the case for the GISAID data base [43] where each sequence is assigned when uploaded, to a single lineage based on which characteristic mutations it carries.

**Figure 9.**
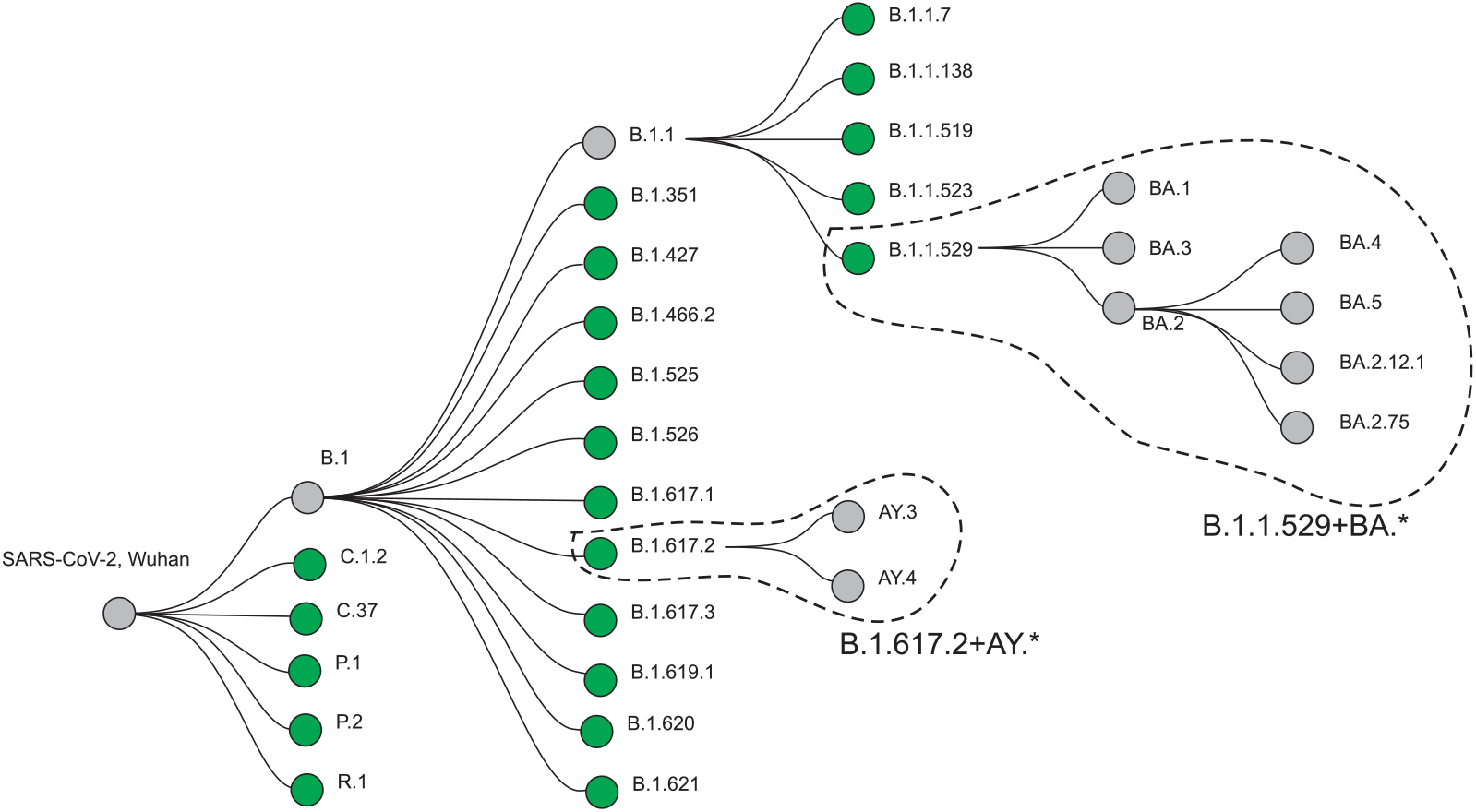
The hierarchical structure of the 28 lineages and sublineages monitored by the BV-BRC SARS-CoV-2 tracking system [5]. This structure reflects their evolutionary trajectories and relationships [22, 24]. The lineages selected for our study (green) are independent in this hierarchy structure.

Our approach takes the hierarchical structure of the 28 LOCs into account by stratifying them. In this study, we only consider one level of this hierarchy, which consists of 21 lineages, with Omicron (B.1.1.529+BA.*) including all BA.* sub-lineages, see Fig. 9. Our choice of these 21 lineages allows us to partition the sequence space into non-overlapping subsets, facilitating the downstream analysis on lineage relative frequency calculations. We follow the characteristic mutation definitions given in Pango Lineages [39] for our considered LOCs.

### Data preparation

SARS-CoV-2 whole genome data was collected from GISAID [43]. All collected sequences were aligned to the reference sequence collected from Wuhan, 2019 (GISAID ID: EPI_ISL_402124). A multiple sequence alignment (MSA) was produced by MAFFT [28]. Each sequence was labeled by collection time, country and classified by lineage.

We the partitioned the sequences into time bins according to their collection times at a weekly resolution, where week 1 equals day 1 to 7, week 2 equals day 8 to 15 and the following weeks are mapped accordingly. We then integrate sequences over three consecutive time bins into one MSA, labeled by the last time bin. In countries like the UK and the US, there were more than 10, 000 sequences per week when the virus is heavily circulated in the human population. In view of computational feasibility, we implemented a sequence cap of 3000 per week via uniform sampling.

The relative frequency for a lineage within the population at a given time was computed based on the lineage classification in the respective time bin. If the sequence was labeled by one of the 21 selected lineages, it was counted as its respective lineage. However if a sequences was labeled by a sub-lineage of one of the 21 selected lineages, then it contributed to the frequency of it parent among the 21 lineages.

### Relational structure construction

For a given MSA, the relational structure is constructed by first selecting polymorphic sites, and then partitioning the polymorphic sites into motifs via clustering.

#### Polymorphic sites selection

Let *M* be an MSA with *m* sequences of length *n*, where each row is a sequence, and each column corresponds to a genetic site. We filter for sites exhibiting sufficient large nucleotide diversity such that a significant co-evolutionary signal may be observed. To this end, we utilize Shannon entropy [42] to identify sites where selection induces evolutionary variation. Namely, *H*(*i*) = −∑ _*x*∈{*A,T,C,G*}_ *p*_*i*_(*x*) log^2^ *p*_*i*_(*x*), where the units of *H* are bits, and *p*_*i*_(*x*) is the probability of the nucleotide *x* appearing in column *i*. Note that there may be gaps in the column of an MSA. If a column contains more than 30% gaps, this column will be filtered out and not considered for further calculations. Otherwise, we ignore the gaps and compute the entropy of the columns based on the remaining nucleotide identities. A site *i* is selected if *H*(*i*) *> h*_*0*_. Here, we set *h*_*0*_ = 0.1.

#### Computing pairwise J-distance between selected sites

We then construct a distance via *joint entropy* and *mutual information* as follows: the *joint entropy H*(*i, j*) of two sites *i* and *j* is defined as

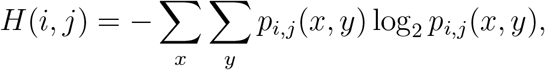

where *p*_*i,j*_ denotes the joint distribution of columns *i* and *j*, i.e., *p*_*i,j*_(*x, y*) specifies the probability of pairs of nucleotides (*x, y*) ∈ {*A, T, C, G*} × {*A, T, C, G*}. Here, we will not consider as a pair *x, y* if either *x* or *y* is a gap. Clearly, the marginal probability distributions for columns *i* and *j* are given by *p*_*i*_(*x*) = ∑_*y*_ *p*_*i,j*_(*x, y*) and *p*_*j*_(*y*) = ∑_*x*_ *p*_*i,j*_(*x, y*), respectively.

The *mutual information I*(*i*; *j*) between sites *i* and *j* is the relative entropy between the joint distribution *p*_*i,j*_(*x, y*) and the product distribution *p*_*i*_(*x*)*p*_*j*_(*y*):

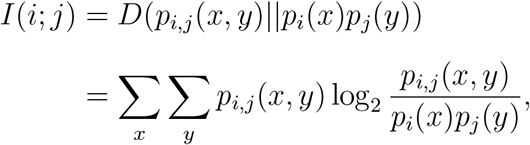

where *D*(*p*||*q*) denotes the *Kullback-Leibler divergence* of the distribution *p* with respect to the distribution *q* [32]. The mutual information *I*(*i*; *j*) quantifies the amount of information shared by two columns *i* and *j*.

Then the *J-distance J*(*i, j*) between two sites *i* and *j* is given by

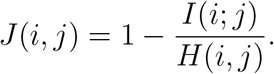

The J-distance represents the information-theoretic counterpart of the Jaccard distance [26, 9]. *J*(*i, j*) satisfies the following properties: 0 ≤ *J*(*i, j*) ≤ 1, *J*(*i, j*) is a pseudo metric, i.e. *J*(*i, i*) = 0, *J*(*i, j*) = *J*(*j, i*) and the triangle inequality *J*(*i, j*) ≤ *J*(*i, k*) + *J*(*k, j*). Furthermore, *J*(*i, j*) is *scale invariant*, that is *J*(*i, j*) is independent of the MSA size, and uniquely determined by the joint distribution of pairs of bases.

#### Constructing motifs via clustering

We utilize highly connected subgraphs (HCS) clustering [23] to compute the relational structure based on the pairwise *J*-distances for the selected pairs of sites. The HCS clustering algorithm is based on the partition of a similarity graph into all its highly connected subgraphs. More precisely, two active columns *v*_*1*_, *v*_*2*_ ∈ *V* are in the same cluster if they belong to the same highly connected subgraph of *G*. HCS clustering does not make any *a priori* assumptions on the number of clusters. Two sites are clustered purely based on their pairwise *J*-distance and it is irrelevant to how many sites are to be clustered. Therefore, the clustering result is robust with respect to the selection of columns with diversity (different choices of *h*_0_ do not impact whether the same two site cluster together).

Based on the pairwise *J*-distance for the selected sites, we construct the *similarity graph G* = (*V, E*) as follows: two vertices *v*_*i*_, *v*_*j*_ ∈ *V* are connected by an edge if their *J*-distance is smaller than the threshold *J*(*i, j*) < *m*_*0*_. In this paper, we set *m*_*0*_ = 0.6. We then group all selected sites into disjoint clusters (motifs) *M*_*1*_, *M*_*2*_, …, *M*_*r*_ via HCS-clustering on the similarity graph. Here, we only consider clusters with size ≥ 5, as most of the currently designated lineages contain at least five characteristic mutations. This yields the relational structure of a given MSA *S* = {*M*_*1*_, *M*_*2*_, …, *M*_*r*_} as collection of clusters.

### Relevancy assessment of a lineage

A lineage *L* is a collection of characteristic mutations, *L* = {*μ*_*1*_, …, *μ*_*k*_}. We will introduce a score, *ϵ*(*L, S*), for assessing the relevancy of *L* by a relational structure *S* = {*M*_*1*_, *M*_*2*_, …, *M*_*r*_}. Let *i* be a site and consider a mapping *Q*(*I*), where *Q*(*i*) = |*M*_*j*_| if *i* ∈ *M*_*j*_ for some 1 ≤ *j* ≤ *r*. Otherwise, *Q*(*i*) = 0.

Let 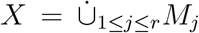 be all sites contained in *S*, and *Y* = {*y*_*1*_, …, *y*_*k*_} where the mutation *μ*_*i*_ takes place at site *y*_*i*_. We then have

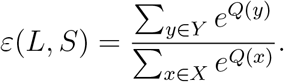

### Alert issuance

Our surveillance system aims to predict whether a lineage *L* will develop into a key lineage for a given country *X*. with respect to *X*, an alert will be issued by our system, regarding lineage *L* at time *t*, if Δ*ε*(*L, S*_*X*_(*t*)) *>* 0.25, where *S*_*X*_(*t*) is the relational structure corresponding to *X* at time *t*.

## Data Availability

The nucleotide sequences of the SARS-CoV-2 genomes used in this analysis are available, upon free registration, from the GISAID database (https://www.gisaid.org/).

## Acknowledgements

We thank Dr. Anindya Dutta and Briana Wilson for the discussion and inspiration for the work. We thank Dr. Andrew Warren for helping us access SARS-CoV-2 data. Dr. Warren pointed out the importance of applying our method to LOC, monophyletic clusters and provided us with numerous references. We thank Mia Shu for helping us to access and process the GISAID data. Many thanks to Maxwell Reidys for his proofreading. This work was partially supported by the VDH Grant PV-BII VDH COVID-19 Modeling Program VDH-21-501-0135.

